# Mutation of the peptide-regulated transcription factor ComR for amidated peptide specificity and heterologous function in *Lactiplantibacillus plantarum* WCFS1

**DOI:** 10.1101/2024.01.25.577207

**Authors:** Michael Brasino, Eli Wagnell, Sila Ozdemir, Srivathsan Ranganathan, Justin Merritt

## Abstract

There is a growing interest in the use of probiotic bacteria as biosensors for the detection of disease. However, there is a lack of bacterial receptors developed for specific disease biomarkers. Here, we have investigated the use of the peptide-regulated transcription factor ComR from S*treptococcus spp*. for specific peptide biomarker detection. ComR exhibits a number of attractive features that are potentially exploitable to create an exquisitely sensitive biomolecular switch for engineered biosensor circuitry within the probiotic organism *Lactiplantibacillus plantarum* WCFS1. By screening a library of ComR mutant protein variants, we identified mutations that increased the specificity of ComR toward an amidated version of its cognate extracellular signaling peptide, demonstrating the potential for ComR to detect this important class of biomarker.

## Introduction

The administration of bacteria as healthcare has been practiced since antiquity and continues today. However, the use of genetically engineered bacteria as diagnostic probes is a relatively recent development. While still a largely untested strategy, there are several advantages to the use of probiotic bacteria for biosensing *in vivo*. For one, multiple bacterial species have been safely consumed by the public for millennia, several of which are also classified as GRAS (Generally Regarded as Safe) by the Food and Drug Administration.^1^ Also, many strains exhibit different tropisms for specific human organs and vary in their proclivity to engraft in human-associated microbiomes, allowing the location and duration of biosensing activities to be tailored by strain selection.^2^ Bacteria have also evolved to detect minute quantities of specific signaling biomolecules, which often yield downstream amplified signals in response. As such, there is a growing interest in exploiting such abilities within genetically engineered bacteria as *in vivo* biosensor probes for a range of diseases.

While bacterial biosensors for markers such as hemin, lactate, hypoxia, and pH have been successfully employed, the ability to reliably detect low-abundance host-derived biomolecules is a far greater genetic engineering challenge and only a small number of successes have been reported.^3–5^ Recently, bacteria have been engineered to detect specific sequences of extracellular human DNA through a combination of CRISPR interference and homologous recombination.^6,7^ Importantly, this detection scheme could be reprogrammed for a range of sequences. Similarly, reprogrammable receptors for specific extracellular biomarkers have been made by fusing nanobodies and other analyte binding motifs to the extracellular domain of the receptor protein kinase CadC in *E. coli*, allowing for the detection of biomarkers in clinical samples.^8,9^ Analogous platforms for specific and reprogrammable receptors will greatly enhance the utility of bacterial biosensors.

Extracellular human peptides are an important class of biomarkers for which bacterial detection schemes have not been reported. For example, gastrin-releasing peptide (GLP) is released by nerves within the stomach to induce gastrin release from G-cells but is also ectopically secreted by many neuroendocrine tumors, such as small-cell lung cancer. Its quantification in serum may be diagnostic and prognostic.^10^ Similarly, capped peptides, which are both amidated at their C-termini and contain pyroglutamylation at their N-termini, are a ubiquitous and understudied class of biomarkers.^11^ Here, we have investigated the bacterial detection of such modified peptides by re-engineering the streptococcal peptide-binding transcription activator ComR.

ComR is a transcription factor that is allosterically regulated by a short peptide secreted by the same bacterial strain as a method of quorum sensing. The import and subsequent binding of this peptide to ComR induces a conformational change that allows ComR to bind a specific DNA consensus sequence, stimulating the transcription of multiple genes, including the *comX* gene, encoding a natural competence-specific sigma factor. As such, this short peptide has been termed *comX*-inducing peptide or XIP, which is encoded by the *comS* gene and is itself regulated by ComR. Both the ComR and XIP sequences vary between species and the ComR from each species recognizes its cognate XIP with varying degrees of specificity.^12^ The availability of this natural sequence vs specificity information has led to several efforts to re-engineer the specificity of ComR for alternative peptide ligands.^13,14^ Importantly, the structure of ComR from *S. thermophilus* has been solved both in complex with its XIP and alone, informing the design of point mutants that have specificities to other species of XIP peptide.^13^ However, the use of ComR to detect non-XIP peptides has not been thoroughly explored.

Engineering ComR to detect human peptides could form the basis for new biosensors in the synthetic biology field. Here, we have investigated the function of ComR from *S. thermophilus* in both a laboratory strain of *Escherichia coli* and in the well-characterized probiotic bacterium *Lactiplantibacillus plantarum* WCFS1. Furthermore, we have taken an initial step toward detecting human biomarker peptides by mutating ComR to specifically bind an amidated peptide.

## Results

### ComR activation by exogenous XIP drives reporter gene expression in *L. plantarum*

We first set out to investigate if ComR could be used to drive gene expression in response to exogenous XIP peptide (LPYFAGCL) in the probiotic strain *Lactiplantibacillus plantarum* WCFS1 (*L. plantarum*). To do this, we inserted the synthetic gene circuit shown in **Figure 1** into the *L. plantarum* genome using a phage recombinase-based method, such that ComR formed a transcriptional fusion downstream of the constitutively expressed ribosomal subunit protein S21 (*rpsU*). In this construct, a heterologous ribosomal binding site (RBS) drives the translation of ComR, which in turn regulates a downstream promoter (P) controlling expression of a secreted nanoluciferase (nLuc), while read-through transcription from ComR to nLuc is prevented by a rho-independent transcriptional terminator (T). Initially, we used a strong RBS and spacer region sequence from *Streptococcus sanguinis* lactate dehydrogenase gene (*ldh*) for the RBS, the *comS* promoter found immediately downstream of ComR in *S. thermophilus* for the promoter, and the widely-used transcriptional termination sequence from *Escherichia coli’s rrnC* gene for the terminator. The resulting strain (v1) secreted nLuc in response to exogenous XIP peptide in complex media. However, the EC50 of XIP for this strain was 238.3 nM (95% Cl [217.4, 261.1]), which was higher than reported for ComR reporter strains of *S. thermophilus* or similar streptococci.^15–17^ To increase the sensitivity of this system, we replaced the *comS* promoter with that of STER_1655 (also from *S. thermophilus* LMD-9, accession ABJ66800.1) as it was previously reported that the STER_1655 reached even higher levels of transcription in response to XIP than ComS.^16^ However, this new strain proved difficult to characterize due to a slow growth rate on agar plates or liquid media. We suspected that despite our inclusion of a transcriptional terminator, read-through transcription from *rpsU* and ComR could still be causing leaky nLuc expression. To further prevent this, we replaced the terminator with a synthetic termination sequence (L3S2P21) which was shown to be stronger than that of *rrnC* in *E. coli* (382.13 AU vs 110.63 AU), and successfully inserted this construct into the *L. plantarum* genome to produce v2.^18^ This strain produced 10x less basal nLuc activity in the absence of XIP than v1, and showed enhanced sensitivity to exogenous XIP, with an EC50 of 79.96 nM (95% Cl [71.36, 89.51]) and a dynamic range of nearly two orders of magnitude. Next, we hypothesized that an overabundance of ComR might be activating nLuc expression without XIP, as ComR has been shown to act as a transcriptional activator (as well as a repressor). We attempted to controllably lower the expression of ComR by first removing the spacer sequence from the RBS (v3) or removing the RBS and spacer entirely (v4). These changes were predicted to lower the relative translation rate from 209.03 to 18.72 and 2.72 AU, respectively. ^19^ While both new strains produced lower levels of nLuc activity in the absence of XIP, they also failed to respond to all but the highest concentrations of XIP tested.

**Figure 1:**
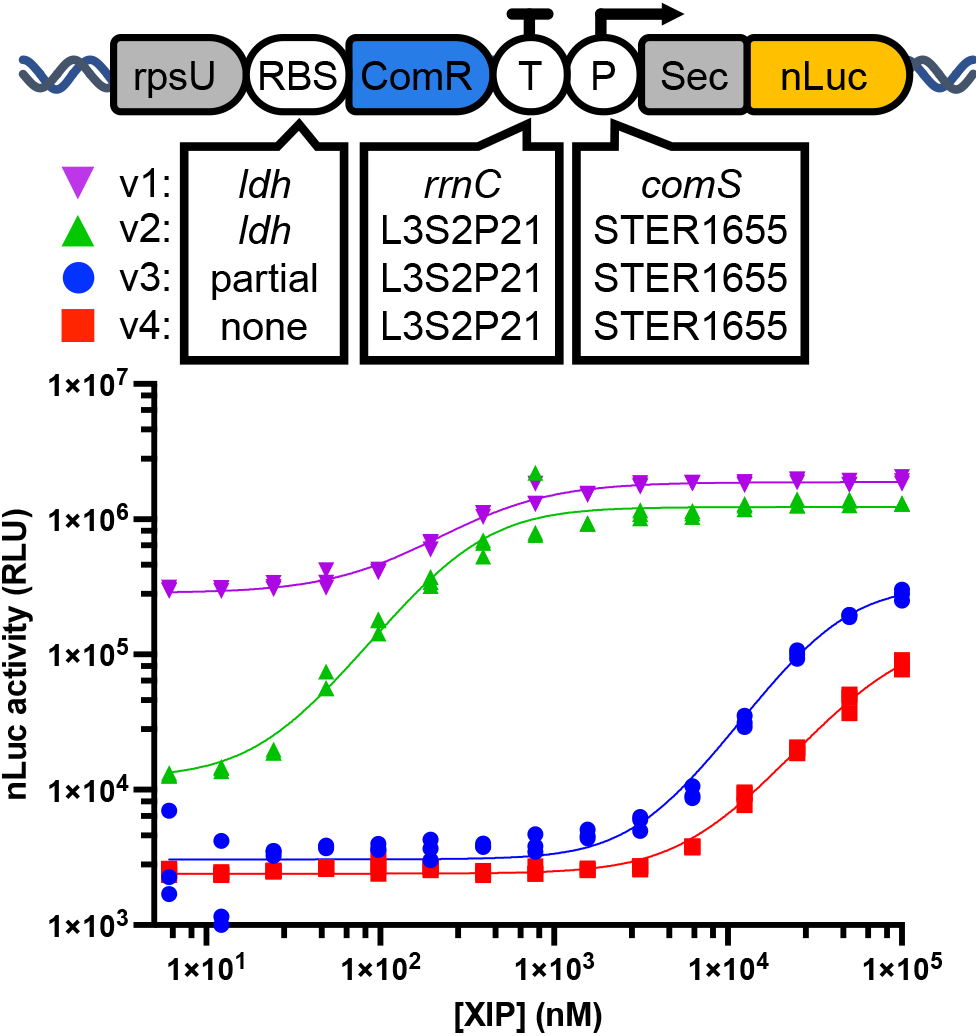
At top is a diagram of the ComR-based genetic circuit inserted in the *L. plantarum* WCFS1 genome behind the endogenous *rpsU* gene. The ribosomal binding site (RBS), transcriptional terminator (T) and promoter sequence (P) for secreted nanoluciferase (nLuc) were modified as shown to create four different versions (v1-4) of this construct. *L. plantarum* carrying each version was grown to mid-logarithmic phase and then transferred to complete cell media (DMEM with 10% FBS) with varying concentrations of XIP peptide. Incubation with each XIP concentration was performed in triplicate (n=3). Nanoluciferase activity after 3 hours of incubation is shown for each replicate. Non-linear regression curves were calculated using a variable slope dose-response model and are shown in the same color for each data set.

### ComR with altered XIP specificity isolated in *E. coli*

We then used the v2 construct as the basis for a ComR mutagenesis and screening system in *E. coli* (shown in **Figure 2)**. Here, ComR was expressed from the low copy number pACYC184 plasmid, with the ComR coding sequence followed downstream by the L3S2P21 synthetic terminator and STER_p1655 promoter. For screening purposes, we replaced the nLuc gene with GFP as a more conveniently read reporter. We also attempted to construct three versions of this screening plasmid, each with a different constitutive promoter driving ComR expression. These were J23119, J23117, and J23116 from the Anderson promoter series of which J23119 was reported to be the strongest and J23116 the weakest. Each promoter was separated from the ComR CDS by a 5’-UTR containing the strong ribosomal binding site (aaagaggagaaa). We failed to produce any colonies from the transformation of the J23119 construct, suggesting that such a high level of ComR expression is prohibitively toxic to *E. coli*. Colonies from both the J23116 and J23117 transformations were grown and tested for GFP expression when grown with or without 10µM XIP peptide. Of these, a single colony from the J23117 transformation showed slightly higher GFP expression with XIP. When sequenced, this construct was found to have a single base pair mutation within the -10 region of the otherwise expected J23117 promoter sequence which was predicted to increase expression (8803.5 AU vs 8513.14 AU).^20^ As this was the best promoter tested, this construct was used for further ComR mutant construction and screening.

**Figure 2:**
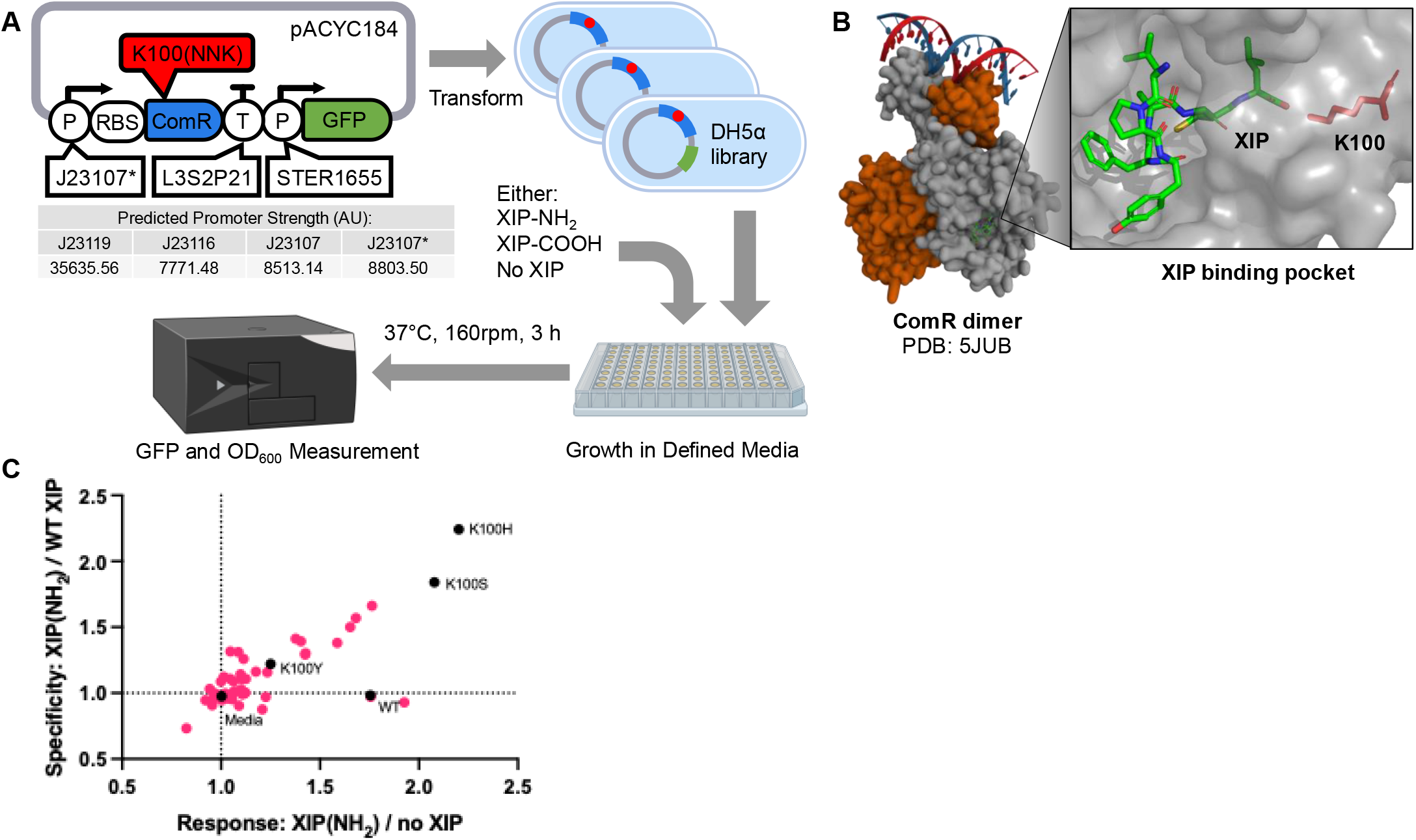
(A) Diagram of the strategy used to screen ComR mutants for altered XIP specificity in *E. coli*. The *E. coli* expression and screening plasmid is shown along with the relative predicted strengths for promoters that were attempted in place of J23107* that was ultimately used for screening. () Structure of ComR from *S. thermophilus*, dimerized and in complex with its XIP and promoter recognition sequence. At right is a detailed view of XIP binding ComR. The XIP and K100 residue of ComR are displayed as stick representation while the rest of ComR is displayed in grey surface respresentation. **(C)** Relative response and specificity of 48 colonies containing ComR mutants. GFP signal was normalized to OD_600_ to determine the relative GFP expression with 50 µM WT or amidated XIP, or to no XIP at all. Response is defined as the ratio of expression with amidated XIP vs no XIP, and specificity as the ratio of expression with amidated XIP vs WT XIP. Colonies with mutations identified through sequencing are shown in black, along with a colony with un-mutated ComR (WT) and a well without bacteria (Media).

We examined the crystal structure of ComR from *S. thermophilus* bound to its cognate wild-type XIP peptide (shown in **Figure 2A**) and found that both lysine at position 100 and threonine at position 92 formed hydrogen bonds with the carboxyl group at the WT XIP C-terminus. We reasoned that mutating the larger lysine residue might allow for the binding of an amidated XIP terminus, and so we created a small library of ComR mutants using degenerate PCR primers containing NNK at the K100 codon. We circularized and transformed the resulting PCR products into *E. coli*, and picked 48 colonies for screening. These colonies, along with the parent ComR expressing strain were compared for their GFP expression after 3 hours of incubation with 50 µM amidated XIP, WT XIP (OH), or no XIP in peptide-free media. Results are shown in **Figure 2C**. Surprisingly, we observed that the wild-type ComR was equally responsive to amidated XIP as it was to WT XIP. Two colonies, in particular, were observed to significantly favor the amidated version and were sequenced along with a colony that showed less activation with either XIP than the wild type as a control. Sequencing revealed that the colonies that favored amidated XIP were mutated to serine and histidine while the colony with a loss of function was mutated to tyrosine. Molecular modeling of serine and histidine mutations suggested they were uniquely suited to accommodate the amidated XIP within the binding pocket (Figure S1).

### Mutant ComRs retain altered XIP specificity in *L. plantarum*

*E. coli* expressing WT ComR or either the K100H or K100S ComR mutants were further characterized with a range of both amidated (NH_2_) and WT XIP concentrations as shown in **Figure 3A**. This confirmed that WT ComR was equally responsive to amidated and WT XIP with EC50 values of 23.85 µM, 95% CI [21.73 to 26.69] and 27.93 µM, [23.06 to 37.54], respectively. Both ComR mutants showed slightly lower sensitivity to amidated XIP than the WT ComR with K100S having an EC50 of 42.96 µM [33.40 to 65.58]. However, both had no detectable response to WT XIP.

**Figure 3:**
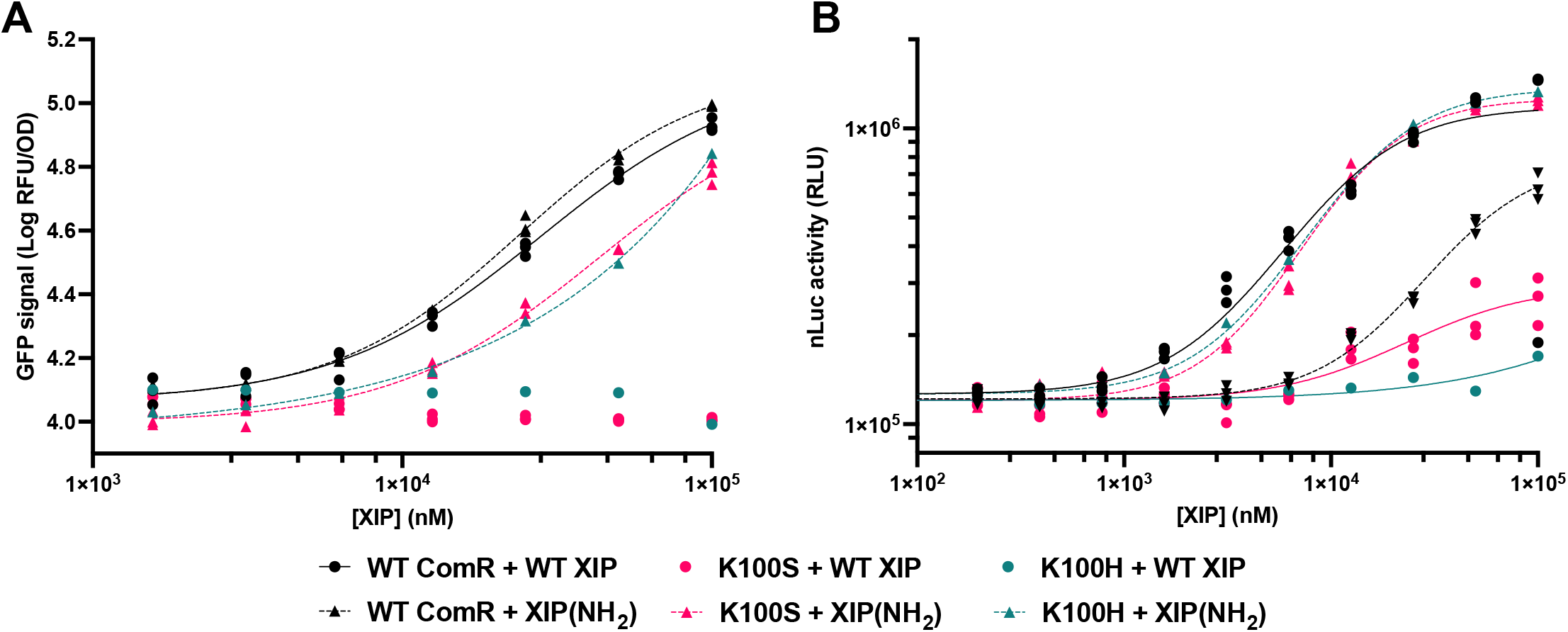
(A) The GFP response to the indicated concentrations of amidated or WT XIP are shown for WT ComR compared to K100S and K100H mutants expressed in *E. coli*. GFP signal after 9 hours of incubation was normalized to OD_600_ and log-transformed. () The nanoluciferase activity after 3 hours of incubation with the indicated concentrations of amidated or WT XIP in complete cell media is shown for WT ComR compared to K100S and K100H mutants expressed in a xylose inducible format on a plasmid in *L. plantarum* WCFS1. For both charts, results of triplicate experiments (n=3) are shown along with variable slope dose-response curves (non-linear regression). The legend for both charts is the same and is shown below.

To determine if this altered specificity was inherent to the ComR sequence regardless of the heterologous host, we then characterized both K100S and K100H mutants in *L. plantarum*. For convenience, this was done on a plasmid expression system. In this plasmid, ComR was driven by a xylose inducible promoter but otherwise contained identical regulatory elements to the v2 circuit integrated in the *L. plantarum* genome earlier. As shown in **Figure 3**, *L. plantarum* was notably less sensitive to WT XIP when expressing ComR from this plasmid than from the genome, with an EC50 of 11.7 µM (95% CI [7.60 to 21.4]). This may have been due to lower expression of ComR from this inducible promoter than when transcribed with the *rpsU* gene. Unlike in *E. coli*, WT ComR in *L. plantarum* showed a preference against amidated XIP, with an EC50 of 48.2 µM (95% CI [40.2 to 65.8]). However, not only did both mutants strongly prefer the amidated XIP, but they both rescued amidated XIP sensitivity to a level similar to WT ComR with WT XIP. Specifically, K100S provided an EC50 of 13.4 µM (95% CI [12.2 to 15.0]) and K100H provided an EC50 of 14.8 µM (95% CI [13.9 to 15.8]).

## Discussion

These results highlight the possibility of using ComR, and peptide pheromone receptors more broadly, for peptide biomarker detection in probiotic bacteria. To realize this possibility, receptors need to be (1) functional in a diverse range of heterologous hosts and (2) easily reprogrammed to detect a range of peptide sequences. As demonstrated here, ComR may meet both conditions, as it can drive gene expression in the distantly related organisms *L. plantarum* and *E. coli*, and it is capable of responding to an amidated variant of the XIP peptide. Further studies will be required to determine the range of peptide substrates that ComR can be adapted to detect. However, our results and previous studies suggest that ComR is highly amenable to reconfiguration of its peptide specificity.^13^

The *E. coli*-mediated selection system developed here might be useful in the re-engineering of ComR. However, it has several notable limitations. Nanomolar concentrations of XIP activated ComR in *L. plantarum*, which is similar to results reported by others in streptococci or *in vitro*. In contrast, micromolar quantities of XIP were required to activate ComR in *E. coli*. It is somewhat surprising that ComR expressed in *E. coli* was capable of responding to XIP at all, as this required exogenous XIP to enter the *E. coli* cytoplasm. We hypothesize that XIP likely transits the *E. coli* cytoplasmic membrane via the native oligo peptide permease (Opp). This is supported by our observation that growth in peptide-free (chemically defined) media was required for ComR to respond to XIP there. The *E. coli* Opp system has been shown to favor shorter peptides up to pentamers, and so the 8 amino-acid long XIP (LPYFAGCL) used here is likely out-competed by the numerous short peptides in rich media.^21^ The effects of the oligopeptide permease specificity in *L. plantarum* may be less prevalent, due to its similarity to *Lactococcus lactis* which readily binds larger peptides.^22^ We attempted to further validate the *E. coli* screening system by cloning a version of *S. thermophilus* ComR containing 5 mutations (R92G, V205A, S248G, S289K, I290T), which was reported to have altered specificity for the 8 amino-acid long XIP from *S. vestibularis* (VPFFMIYY).^13^ This showed no reactivity to XIP from *S. thermophilus* (as expected) but only a slight increase in GFP signal with 100 µM of *S. vestibularis* XIP. This mutant was reported to have a lower affinity, but differences in the binding of the *E. coli* Opp to each XIP sequence could not be ruled out. In the future, this screening system might be improved by co-expressing native or streptococcal OppA proteins with ComR, or by co-expressing the desired peptide ligand in the *E. coli* cytoplasm.

## Methods

### Strains and Routine Culture Conditions

*Lactiplantibacillus plantarum* WCFS1 was purchased from the American Type Culture Collection (BAA-793) and was routinely cultured in MRS (De Man–Rogosa–Sharpe) media purchased from Research Products International (L11000-1000.0) at 37 C under 5% CO_2_ atmosphere. MRS was either autoclaved or filter sterilized using 0.2 µm vacuum filters to prevent discoloration. *Escherichia coli* strain DH5α was purchased from New England Biolabs (C2987H) and routinely cultured in LB media (Sigma) at 37 C with shaking.

### Genetic Modification of *L. plantarum*

ComR-based sensor constructs were inserted into the *L. plantarum* WCFS1 (WCFS1) genome using a previously reported phage-recombinase method.^23^ Genetic sensor constructs were first constructed on the shuttle plasmid pTRKH2 (Addgene Plasmid #71312) and sequence verified. Then, both the sensor construct and erythromycin resistance cassette from pTRKH2 were PCR amplified together, and 1 kb long homology arms were PCR amplified from the WCFS1 genome. After gel extracting all PCR products, the homology arms were appended to the sensor amplicon using overlap extension PCR. This extended product was gel-extracted as well. All WCFS1 transformations were done via electroporation following a previously documented protocol.^24^ WCFS1 was first transformed with helper plasmid pLH01 (Addgene Plasmid #117261) containing phase recombinases under an inducible promoter. This helper strain was prepared for electroporation again with the addition of inducing peptide (Sequence MAGNSSNFIHKIKQIFTHR, produced by Genscript) to induce recombinase expression for approximately 1 hr during the mid-logarithmic growth phase. Transformants were selected on MRS agar plates containing 10 µg/ml erythromycin. Correct insertion of the sensor circuit was confirmed by PCR amplifying the surrounding section of the genome using primers that sat outside of the 1 kb homology arms, and then sequencing the resulting product.

### Characterizing ComR sensor sensitivity in *L. plantarum*

*L. plantarum* strains containing genome-integrated ComR sensors were stored in glycerol stocks at -80 C until needed. These were grown overnight (at least 16 hours) in MRS with 10 µg/ml erythromycin at 37 C with 5% CO_2_ without shaking. They were then diluted to an OD_600_ of approximately 0.3 in fresh media and grown under the same conditions until reaching mid-logarithmic phase (OD_600_ = 0.6 - 1.0). Then, 1 ml of OD_600_ = 0.8 culture (or equivalent OD_600_ x ml) was centrifuged at 4,000 g for 10 minutes. Each cell pellet was resuspended in 100uL of Phosphate Buffered Saline (Fisher #BP399) and 2uL of each resuspended cell culture was added to a 384 well plate (Corning #3764). Then, 18 uL of XIP peptide (sequence LPYFAGCL, Genscript) diluted in DMEM (Gibco# 11995-065) +10% FBS (Cytiva #SH30396.03) was added to the corresponding wells. For each strain and XIP concentration, three replicate wells were used. The plate was then allowed to incubate for 3 hrs at 37 C. To read the resulting nLuc activity, 3 ul was taken from each well and placed into a black flat bottom 96-well plate (Corning #3925). Then 30 µl of premixed Nano-Glo substrate/buffer (Promega #N1138 and Promega #N1128 prepared following manufacturer’s instructions) and 27 µl of diluted 1 M Tris-HCl (pH 8.0) (Invitrogen #15568-025) were simultaneously added to each well. The plate was then immediately read on a Tecan Spark 20M plate reader programmed to shake the plate for 30 seconds, and then measure the luminescence between 430 nm and 500 nM for 1 second.

### Characterization of ComR specificity in *E. coli*

ComR sensor circuits were constructed on plasmid pACYC184 (American Type Culture Collection #37033) such that ComR activation promoted GFP expression. These were verified using NGS whole plasmid sequencing (Plasmidsaurus) and transformed into chemically competent DH5α which were stored as glycerol stocks at -80 C until needed. To test for ComR specificity, transformants containing ComR sensor plasmids were grown overnight in LB broth with 10µg/ml tetracycline (Gold Bio #T-101-25). The next day, bacteria were pelleted by centrifugation at 6800g for 3 mins, then diluted at 1:100 in Hi-Def Azure Media (Fisher Scientific # 3H5000) supplemented with 1% Glucose and various concentrations of peptide in 100 µl in a 96well plate. All peptides were synthesized by Genscript. Bacteria were incubated in a large radius shaker (Eppendorf S441) at 37 C for 9 h, then GFP fluorescence was read using a TECAN plate reader. Parafilm or plastic plate covers were used to prevent cultures from drying.

### Statistical Analysis and Curve Fitting

All statistics and curve fitting were done using the Prism 10 software package for Mac OS.

## Acknowledgements

This project was supported by funding (Full7340220, Full 2021-1439) from the Cancer Early Detection Advanced Research Center at Oregon Health & Science University, Knight Cancer Institute (Project Lead: Michael Brasino). Additional support was awarded from the Collins Medical Trust (ACNCR1235, PI: Michael Brasino).

## Notes

Conflict of Interest: The authors declare no conflict of interest.

